# Saprotrophic soil fungus dissolves lithium lepidolite through inositol rescue metabolism

**DOI:** 10.64898/2026.07.28.741394

**Authors:** Sandra LaBonte, Cayden Perdue, Thomas Wietsma, Vanessa Paurus, Young-Mo Kim, Hsin-Mei Kao, Zihua Zhu, Erin Nuccio, Jennifer Pett-Ridge, Mary Lipton, Arunima Bhattacharjee

## Abstract

Microbial weathering of minerals represents a key biogeochemical process through which microorganisms access essential nutrients locked within rock and ore substrates. In this study, we investigated the metabolic response of a model fungus during growth on lithium (Li) ore to elucidate the mechanisms underpinning biologically mediated mineral weathering and subsequent metal mobilization. Our results revealed a distinct metabolic shift in the fungus when cultivated on Li ore, characterized by altered patterns of organic acid production and energy metabolism. This shift coincided with measurable weathering of the Li ore matrix and the consequent release of soluble Li into the surrounding environment, demonstrating a direct link between fungal metabolic reprogramming and mineral breakdown. Importantly, these observations are consistent with metabolic shifts documented in previous studies of mineral weathering by this fungus, suggesting that such responses constitute a conserved and reproducible strategy employed during the colonization of mineral substrates. Although the present work was conducted using a single model microorganism under controlled conditions, the findings carry broader ecological implications. Natural microbial communities inhabiting mineral-rich environments may undergo analogous metabolic shifts when weathering minerals to acquire limiting nutrients, thereby contributing to large-scale elemental cycling and metal release. Understanding these processes not only advances fundamental knowledge of microbe–mineral interactions but also informs emerging applications in biomining and bio-based recovery of critical metals such as lithium. Collectively, this study highlights the central role of microbial metabolism in driving mineral weathering and offers a framework for predicting and harnessing similar processes within complex microbial communities.

**IMPORTANCE:** The model fungus exhibits a distinct, reproducible shift in metabolism—particularly in organic acid production and energy pathways—when grown on Li ore, directly linking cellular metabolism to mineral breakdown. This metabolic shift promotes weathering of the ore matrix and the release of soluble lithium, demonstrating a biological route for liberating a critical metal from its mineral host. Because similar shifts occur across different minerals, natural microbial communities likely employ comparable strategies to weather minerals and acquire limiting nutrients, contributing to global elemental cycling. These findings provide a foundation for sustainable biomining and bio-based recovery of lithium and other critical metals, offering a lower-impact alternative to conventional extraction methods.

## INTRODUCTION

Microorganisms in soils, the rhizosphere, and the subsurface routinely dissolve minerals to access nutrients – particularly potassium, phosphorous, iron, and trace metals. This biologically driven weathering proceeds through the concerted action of proton attack (acidolysis), organic ligand complexation (complexolysis), redox transformation (redoxolysis), and physical disruption of mineral surfaces (1–4). While the geochemical consequences of microbial weathering — element fluxes, secondary mineral formation, soil development — have been studied extensively, the metabolism of the organism doing the weathering is understudied.

The limited evidence available suggests that microbial metabolism reorganizes in response to resource availability and the chemical stresses imposed by the dissolution of mineral weathering. Across diverse mineral systems, responses include increased exploitation of inorganic redox chemistry on mineral surfaces(2, 5), elevated production of organic acids, siderophores and other chelators(6–8), and extracellular polymeric substances (EPS), that can acidify and complex mineral surfaces and enhance dissolution(9, 10). In heterotrophic bacteria and fungi, production of low-molecular-weight organic acids — citric, oxalic, gluconic, succinic, and malic acids — increases during growth on mineral substrates relative to mineral-free controls, and this “overflow” acid secretion has been interpreted as a nutrient-acquisition strategy analogous to siderophore production under iron limitation (4, 6–8, 11–13). However, mineral dissolution simultaneously releases potentially toxic cations and anions into the microenvironment, imposing ionic, osmotic, and oxidative stresses whose metabolic consequences extend far beyond organic acid production. How central carbon metabolism, redox defense, osmolyte biosynthesis, amino acid allocation, and secondary metabolite production are coordinately reorganized during active mineral weathering — and whether the resulting metabolic state feeds back on dissolution kinetics — remains largely uncharacterized. Most studies of microbial mineral dissolution have measured extraction efficiencies and organic acid concentrations in spent medium without examining how the organism’s global metabolic program is restructured to sustain mineral attack and to cope with its chemical consequences (14–17).

Filamentous fungi are particularly effective agents of mineral weathering. Their branching hyphal networks maintain prolonged, intimate contact with mineral grains, and their metabolism generates a diverse arsenal of organic acids and chelating metabolites. These compounds create steep, localized gradients of low pH and high ligand activity at the hypha–mineral interface (1, 18–20). These same properties have motivated growing interest in fungi as agents of biomining — biological extraction of critical elements from ores and mine wastes (21–23). Among biomining targets, lepidolite [KLi₂Al(Si₄O₁₀)(F,OH)₂] is an especially informative model system for studying metabolic responses to mineral weathering. It is one of the most abundant lithium-bearing minerals in pegmatite deposits, yet its refractory crystal chemistry renders it largely uneconomical for conventional extraction, and it is frequently discarded as waste during spodumene-focused mining operations (24–26). Crucially for mechanistic studies, lepidolite contains negligible structural iron (27, 28), effectively excluding redox-mediated dissolution and isolating the metabolic contributions of acid- and ligand-promoted attack. At the same time, progressive dissolution releases Li⁺, Al³⁺, and F⁻ — a combination of ionic stresses whose effects on fungal metabolism have not been examined (17, 24). Of these, Li⁺ is uniquely tractable because it has a specific, well-characterized intracellular enzyme target: inositol monophosphatase (IMPase), which catalyzes the recycling of myo-inositol from inositol phosphates and the final step of de novo inositol biosynthesis (29). In *Saccharomyces cerevisiae*, Li⁺ inhibition of IMPase triggers a conserved rescue response — transcriptional induction of INO1 (myo-inositol-1-phosphate synthase) — that synthesizes myo-inositol de novo from glucose-6-phosphate (30–32). Because glucose-6-phosphate is the shared entry substrate for glycolysis, the pentose phosphate pathway, and INO1-mediated inositol biosynthesis, this rescue response has the potential to restructure the entire carbon economy of the cell — with downstream consequences for organic acid production, redox balance, and osmolyte biosynthesis that could, in principle, feedback on the mineral surface. Whether this cascade operates during fungal mineral weathering has not been tested.

Here, we investigate how the filamentous fungus *Fusarium* sp. DS 682, isolated from the rhizosphere of *Bouteloua gracilis* at the Konza Prairie Biological Station in Kansas (33), reorganizes its metabolism during lepidolite dissolution. We quantify lithium mobilization and partitioning between fungal biomass and medium by inductively coupled plasma mass spectrometry (ICP-MS). We also map lithium association with hyphae by time-of-flight secondary ion mass spectrometry (ToF-SIMS), and profile secreted metabolites by gas chromatography– mass spectrometry (GC-MS) to connect dissolution chemistry to the underlying metabolic program. We show that lithium bioleaching by *Fusarium* sp. DS 682 does not arise from a specialized mineral-weathering strategy. Instead, Li⁺ liberated from the dissolving mineral inhibits the inositol recycling pathway, triggering a compensatory rescue response that diverts glucose-6-phosphate toward de novo inositol biosynthesis and the pentose phosphate pathway at the expense of glycolysis and extracellular glucose oxidation. The resulting carbon reallocation — constrained glycolytic input to an active but partially bottlenecked TCA cycle, selective upregulation of oxalate biosynthesis, enhanced NADPH production for glutathione-based redox defense, and polyol osmolyte diversification — produces the organic acid overflow that drives further mineral dissolution, closing a self-reinforcing feedback loop in which a specific enzyme–ion interaction propagates through central carbon metabolism to reshape the organism’s geochemical impact on the mineral surface.

## RESULTS

### Fusarium sp. DS682 Grows on Lepidolite and Accumulates Lithium

On low-glucose agar (0.4% w/v glucose, 0.1% w/v yeast extract, 0.1 M NaCl; hereafter SA), *Fusarium* sp. DS 682 developed as a thin, sparse mycelial film with fine, filamentous hyphae (Fig. 1A and Supplementary Fig. 1A). In contrast, when cultured on the same media supplemented with 10% (w/v) lepidolite (LiA), the fungus produced markedly denser, bundled hyphae (Supplementary Fig 1B). Similar hyphal aggregation has been observed in fungi colonizing metal-bearing mineral substrates (1, 18, 34) and may reflect both a protective response to lithium toxicity and the development of transport networks for nutrient transport (34–36). Lepidolite-grown cultures also exhibited visibly enhanced purple pigmentation compared to sugar agar controls (Fig. 1A); notably, faint purple coloration was also observed on SA plates, indicating that pigment production is not exclusive to mineral contact but is intensified by it. This pigmentation is consistent with stress-induced activation of secondary metabolite biosynthesis. ToF-SIMS imaging of hyphae grown on lepidolite-amended medium revealed lithium signal distributed throughout the mycelial network (Fig. 1B, Supplementary Fig. 2), providing direct spatial evidence that *Fusarium* sp. DS 682 is capable of solubilizing lepidolite and accumulating lithium within its biomass.

**Figure 1.**
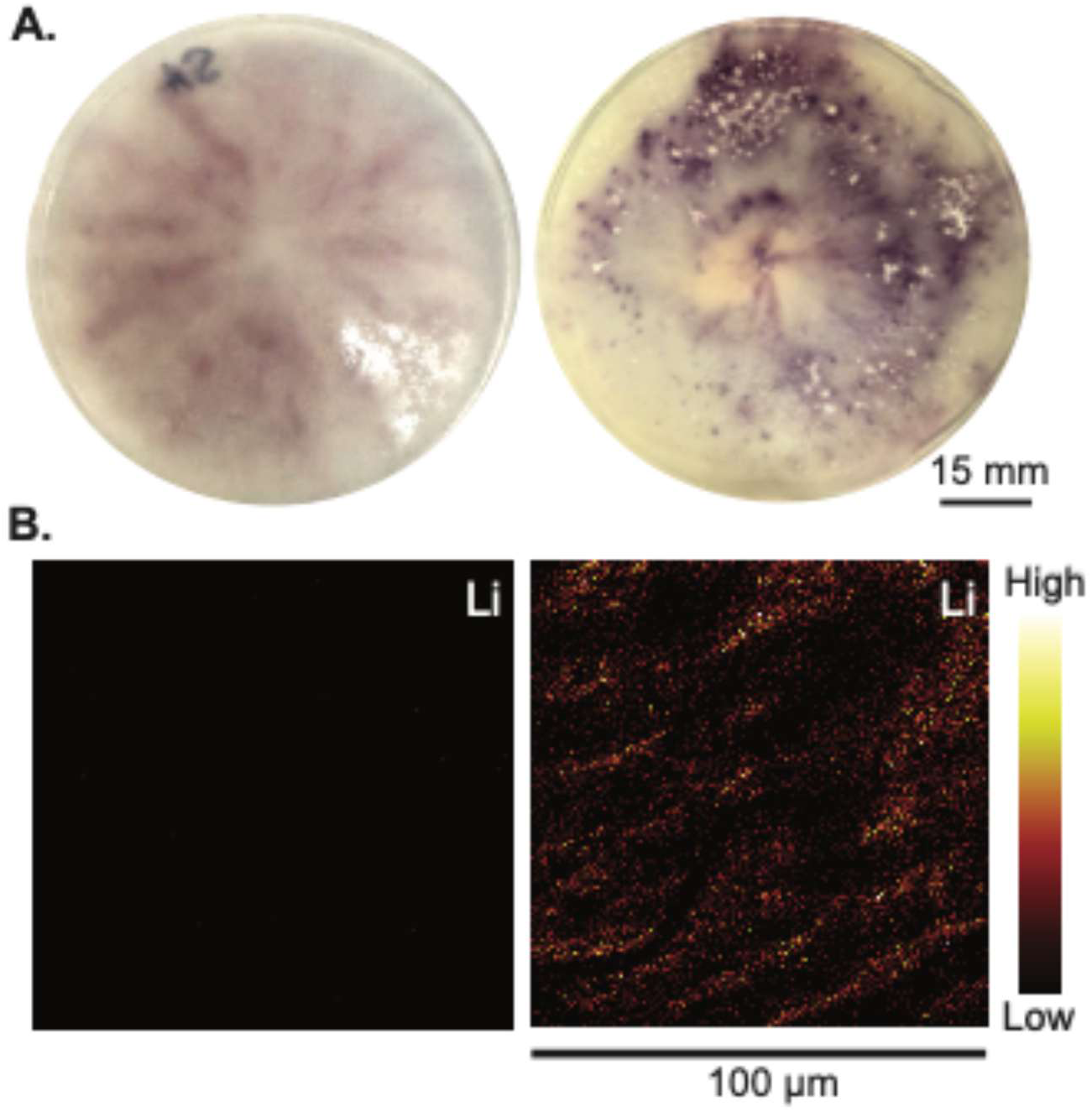
Fusarium sp. DS 682 colony morphology and lithium distribution on sugar agar and lepidolite-amended medium. **(A)** Representative photographs of Fusarium sp. DS 682 colonies grown for six weeks at 25°C on sugar agar (left) or sugar agar supplemented with 10% (w/v) lepidolite (right) **(B)** ToF-SIMS ion maps showing the spatial distribution of Li⁺ signal on mycelium harvested from sugar agar (left) and lepidolite-amended medium (right). Signal intensity is displayed on a thermal color scale from low (black) to high (bright yellow/white). Scale bar, 100 µm (applies to both panels in B).

### *Fusarium* sp. DS 682 Selectively Mobilizes Lithium from Lepidolite

To evaluate the capacity of *Fusarium* sp. DS 682 to dissolve lepidolite and the fate of the liberated ions, concentrations of Li and Fe were quantified by ICP-MS in both the fungal biomass (Fig. 2A, Table S1) and the underlying agar medium (Fig. 2B, Table S1). Iron was measured alongside lithium because in many critical mineral deposits, target elements are structurally associated with Fe-bearing phases whose dissolution is a prerequisite for element liberation (2). Iron concentrations in the mycelium remained at baseline levels regardless of lepidolite exposure (Fig. 2), consistent with the inherently low Fe content of the lepidolite (28). In contrast, Li concentrations in the biomass of fungi grown on lepidolite-amended medium were approximately 12-fold higher than in control mycelium, providing direct evidence of lithium uptake and bioaccumulation by the fungus. Analysis of the agar medium revealed a parallel trend: Li concentrations in agar underlying fungal cultures grown with lepidolite were markedly elevated — approximately 1.8-fold higher — relative to uninoculated lepidolite-containing controls, while Fe levels remained largely unchanged (Fig. 2B). These results demonstrate that *Fusarium* sp. DS 682 actively solubilizes lithium from lepidolite through acid- or ligand-promoted dissolution of the aluminosilicate framework rather than through Fe redox cycling, and that mobilized Li⁺ partitions into both the fungal biomass and the surrounding medium.

**Figure 2.**
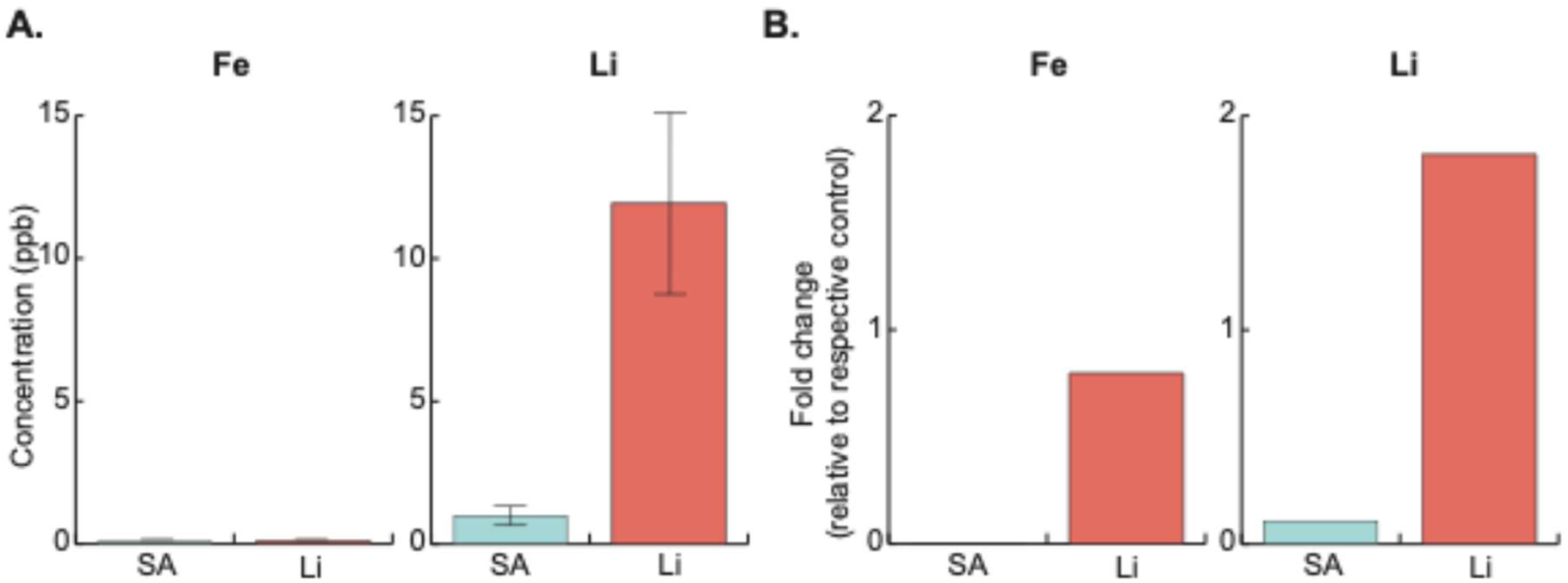
Fungal uptake and release of lithium ions from lepidolite. Concentrations of Fe and Li were quantified by ICP-MS in the fungal biomass **(A)** and the underlying agar medium **(B)**. In both panels, the control (blue) represents fungi grown on sugar agar without lepidolite, and the treated sample (pink) represents fungi grown on lepidolite-amended agar. **(A)** Fe and Li concentrations measured in ground fungal biomass. **(B)** Fe and Li concentrations measured in the agar medium, normalized to the respective uninoculated agar controls (sugar agar for controls, lepidolite agar for treated samples).

### Lithium Stress Redirects Central Carbon Metabolism Toward Inositol Rescue, the Pentose Phosphate Pathway, and Organic Acid Overflow

To investigate the metabolic mechanisms underlying lepidolite weathering and uptake, untargeted GC-MS-based metabolite profiling was performed on fungal cultures grown on sugar agar (SA) and lepidolite-amended agar (LiA), each compared with their respective uninoculated controls. A total of 57 metabolites were identified across all conditions (Table S2). The largest fold-changes on lepidolite-amended medium relative to sugar agar were centered on organic acid production, osmolyte accumulation, and stress defense. Below, differentially abundant metabolites are organized by functional category; compounds exhibiting modest changes are reported in Table S2.

*Myo-inositol elevation and glucose-6-phosphate competition.* Myo-inositol, the central precursor for phosphoinositide signaling and phosphatidylinositol membrane biosynthesis, was approximately 4-fold elevated in LiA fungal cultures relative to SA cultures (Fig. 3A, Supplementary Fig. 3C). This accumulation is consistent with compensatory upregulation of myo-inositol-1-phosphate synthase (INO1), which catalyzes de novo inositol biosynthesis from glucose-6-phosphate — a well-characterized rescue response to Li⁺ inhibition of inositol monophosphatase in *Saccharomyces cerevisiae* (31, 32). Because INO1, the oxidative pentose phosphate pathway (PPP), and glycolysis all compete for glucose-6-phosphate, concurrent engagement of inositol rescue and PPP flux is predicted to constrain glycolytic throughput.

**Figure 3.**
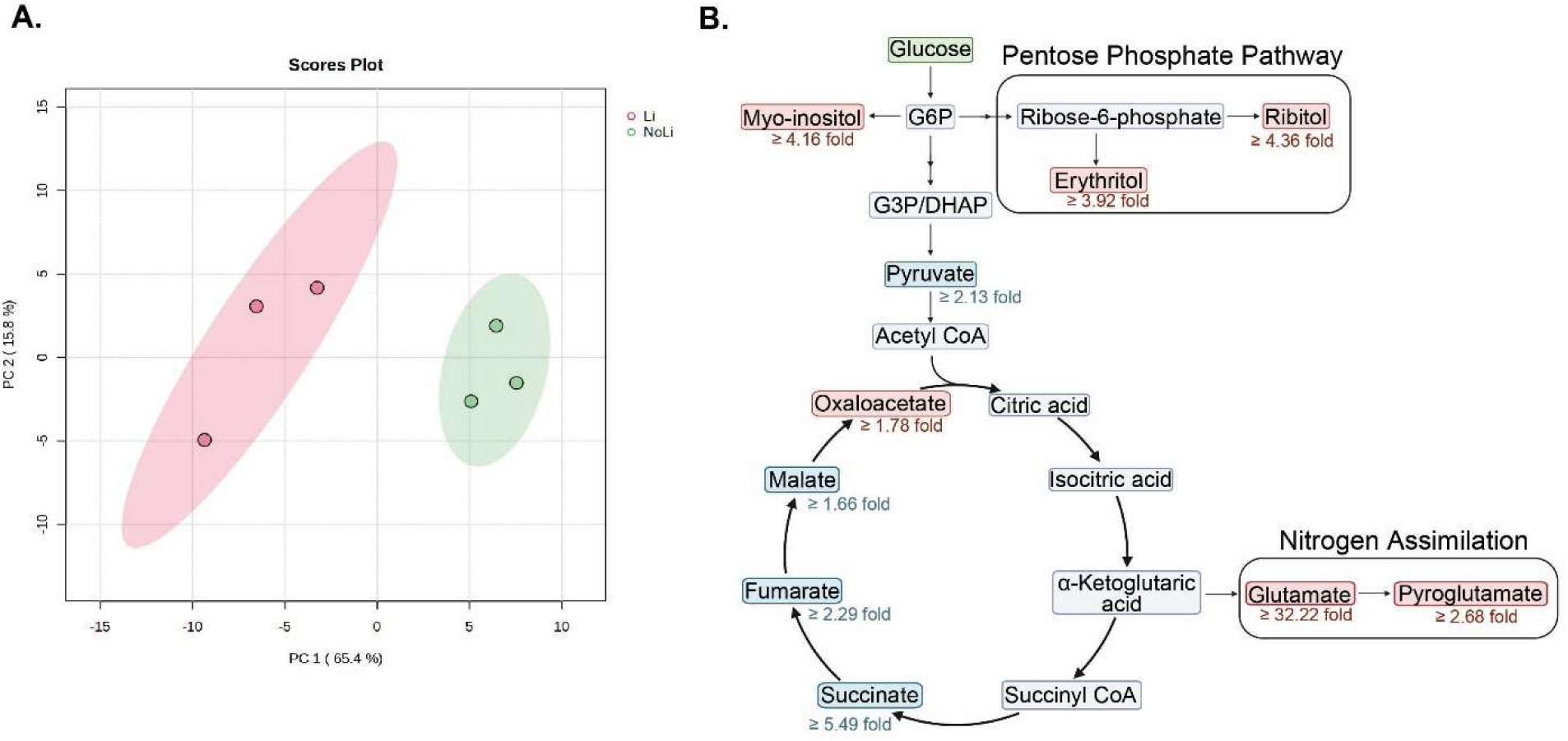
Metabolomic profiling of fungi grown with and without lepidolite. **(A)** Principal component analysis (PCA) scores plot comparing the metabolomic profiles of fungi grown on lepidolite-amended agar (Li, pink) and sugar agar without lepidolite (NoLi, green). PC1 and PC2 explain 65.4% and 15.8% of the total variance, respectively. **(B)** Differentially abundant metabolites mapped onto central carbon and nitrogen metabolism pathways, including glycolysis, the TCA cycle, the pentose phosphate pathway, and nitrogen assimilation. Red fold-change values indicate metabolite increases in the lepidolite condition relative to the no-lepidolite control. Metabolites highlighted in red boxes were significantly elevated in the lepidolite-grown fungi. Metabolites highlighted in blue boxes were significantly elevated in the fungi grown on sugar agar.

*Pentose phosphate pathway engagement.* Consistent with this prediction, D-ribitol (derived from ribose/ribulose intermediates) was more than 4-fold elevated and erythritol (derived from erythrose-4-phosphate) more than 3-fold elevated in LiA cultures relative to SA cultures (Fig. 3B). Both sugar alcohols are products of PPP carbon. Pyruvate, the terminal glycolytic product, was lower in LiA extracts than in SA extracts despite active TCA metabolism under both conditions, a pattern consistent with reduced glycolytic flux when glucose-6-phosphate is diverted toward INO1 and the PPP (Supplementary Fig. 3A&B).

*Gluconate depletion.* D-Gluconic acid, normally produced by extracellular glucose oxidase activity, was markedly depleted in LiA cultures compared to all other conditions (Fig. 3A, Supplementary Fig. 3B, Table S2). This suggests that under lithium stress, glucose is preferentially imported and phosphorylated to glucose-6-phosphate — feeding the elevated intracellular demand from INO1 and the PPP — rather than being oxidized extracellularly to gluconate.

*TCA cycle intermediates and organic acid overflow.* Both SA and LiA cultures showed elevated TCA cycle intermediates relative to their respective uninoculated controls (Supplementary Fig. 3A). On SA, pyruvate (∼2-fold), succinate (∼5-fold), and fumarate (∼2-fold) were most prominently elevated, with absolute abundances of most TCA intermediates higher on SA than on LiA. On LiA, citrate, α-ketoglutarate, succinate, fumarate, and malate were all elevated relative to uninoculated LiA controls, though at lower absolute levels than on SA — a pattern consistent with reduced glycolytic input (less pyruvate entering the TCA cycle) rather than reduced TCA enzyme activity. Citric acid was comparably elevated (∼3-fold over uninoculated controls) under both growth conditions.

*Selective oxalate upregulation.* Among organic acids implicated in mineral dissolution, oxalic acid was the only species selectively elevated in LiA cultures relative to both uninoculated controls and SA cultures (Fig. 3A, Supplementary Fig. 3A). Oxalate is synthesized via oxidative cleavage of oxaloacetate (oxaloacetate hydrolase) or through the glyoxylate cycle (6, 37). Glycolic acid, a metabolic precursor of glyoxylate, was also modestly elevated under lithium exposure, consistent with glyoxylate pathway engagement.

### Lithium Exposure Activates Oxidative Stress Defense and Reconfigures Amino Acid and Lipid Metabolism

*Glutathione cycling and oxidative stress markers.* L-Pyroglutamic acid, a product of glutathione turnover via the γ-glutamyl cycle, was substantially elevated in LiA fungal cultures relative to both uninoculated LiA controls and SA cultures (Fig. 3C, Supplementary Fig. 3C). L-Glutamic acid, the central substrate for glutathione biosynthesis, showed the largest fold-change of any detected metabolite, with approximately 32-fold higher levels in LiA cultures relative to sugar agar cultures (Fig. 3C). L-Serine, which feeds into cysteine biosynthesis via the transsulfuration pathway — providing the rate-limiting substrate for glutathione production — was also selectively elevated under lithium exposure (38, 39). 2-Hydroxyglutaric acid, a marker of mitochondrial redox imbalance formed via reduction of α-ketoglutarate under high NADH/NAD⁺ ratios(40), was elevated relative to controls under both SA and LiA conditions but co-occurred with expanded glutamate and pyroglutamate pools specifically on LiA.

*Amino acid reallocation.* Branched-chain amino acid (BCAA) pathway intermediates were detected under both conditions but showed distinct directionality (Supplementary Fig. 3D). On LiA, the biosynthetic intermediate 2,3-dihydroxyisovalerate (valine pathway) was selectively elevated. On SA, catabolic intermediates — 3-hydroxyisovaleric acid, 3-hydroxyisobutyric acid, and α-ketoisovalerate — were more prominent, indicating a shift from BCAA catabolism (energy recovery on SA) to BCAA biosynthesis (protein replacement/stress support on LiA).

*Membrane lipid suppression.* SA cultures were distinguished by elevated ergosterol, oleic acid (C18:1), linoleic acid (C18:2), stearic acid (C18:0), capric acid (C10:0), and behenic acid (C22:0) relative to LiA cultures or uninoculated controls (Table S2). This lipid-rich profile was not observed under lithium exposure, where carbon appeared directed toward stress-protective pathways rather than membrane biogenesis.

*Compatible solute diversification.* Glycerol and trehalose were elevated under both SA and LiA conditions relative to uninoculated controls, consistent with baseline osmolyte production. However, the osmolyte pool on LiA was qualitatively expanded to include substantial erythritol and D-ribitol — PPP-derived polyols not prominent on SA (Fig. 3B). This diversification suggests that ionic stress from Li⁺ requires a broader compatible solute spectrum than osmotic stress alone.

Metabolites showing minimal or non-significant differences across conditions — including lactic acid, glyoxylic acid, benzoic acid, 5-hydroxymethylfurfural, 2-hydroxypyridine, maleimide, 2,4-dihydroxybutanoic acid, 3,4-dihydroxybutanoic acid, L-threonic acid, and arachidic acid — are reported in Table S2.

## DISCUSSION

### Lithium-Induced Hyphal Bundling Reflects Both Mineral-Stress Cord Formation

While the capacity of filamentous fungi to solubilize lithium from mineral substrates is increasingly recognized (15, 16, 25, 41), the metabolic and cellular mechanisms driving fungal lithium bioleaching remain poorly understood. The results presented here reveal that lithium dissolution by *Fusarium* sp. DS 682 is not driven by a specialized weathering strategy but instead emerges from an integrated stress response in which morphological reorganization, organic acid overflow, osmolyte accumulation, and secondary metabolite overproduction are mechanistically coupled to mineral dissolution. Central to this response is a specific enzyme-ion interaction: Li liberated from the dissolving mineral inhibits inositol monophosphatase (IMPase), triggering a conserved INO1-dependent rescue response that diverts glucose-6-phosphate away from glycolysis and toward de novo inositol biosynthesis and the pentose phosphate pathway (PPP). This single perturbation propagates through central carbon metabolism to produce the organic acid, polyol, and redox-defense signatures described below.

The reorganization of *Fusarium* sp. DS 682 mycelium into dense hyphal bundles on lepidolite-amended medium is morphologically consistent with cord formation, a well-characterized response of filamentous fungi to toxic metal-bearing mineral substrates . Hyphal cords— organized, parallel aggregates of hyphae—serve dual functions in fungi colonizing challenging substrates: they reduce the surface-area-to-volume ratio exposed to toxic ions, conferring protection(1, 18), and they establish long-distance translocation networks for bidirectional transport of water, nutrients, and solubilized elements (34–36). Fomina et al. (2005) reported analogous hyphal aggregation and cord-like morphologies in *Beauveria caledonica* colonizing toxic metal minerals, linking cord formation to enhanced oxalic acid secretion. From a bioleaching perspective, the functional consequences of bundling may be paradoxically beneficial for mineral dissolution. Dense hyphal cords concentrate organic acid secretion at the mineral interface, creating localized zones of low pH and high chelator concentration that may be more effective per unit biomass than diffuse acid secretion from a thin, expansive mycelial film (1, 18). The ToF-SIMS data, which revealed lithium signal distributed throughout the bundled hyphal network, are consistent with these cords being actively engaged in lithium uptake and transport. Practically, because cords concentrate acid secretion and serve as transport conduits, promoting controlled bundling may improve lithium recovery.

The lepidolite-grown cultures also exhibited visibly enhanced purple pigmentation relative to sugar agar controls, although faint purple coloration was also observed on sugar agar, indicating that pigment production is not strictly mineral-induced but is intensified on lepidolite. We did not chemically characterize the pigment in this study, and so its identity remains to be confirmed but purple pigmentation in *Fusarium* is characteristic of naphthoquinone polyketide pigments such as aurofusarin and the fusarubin derivatives, whose biosynthetic gene clusters are transcriptionally upregulated under oxidative stress and nutrient limitation (42–44). If confirmed, such pigments could serve as extracellular ROS scavengers and metal chelators at the hypha-mineral interface.

### Lithium Solubilization and Selective Dissolution of the Lepidolite Lattice

ICP-MS analysis confirmed that *Fusarium* sp. DS 682 solubilizes lithium from lepidolite and accumulates the mobilized Li⁺ within its biomass, with an approximately 12-fold increase in mycelial lithium concentration on lepidolite-amended medium relative to controls. Concurrently, lithium concentrations in the underlying agar were markedly elevated in the presence of the fungus relative to uninoculated lepidolite controls, indicating that a substantial fraction of solubilized lithium partitions into the extracellular environment.

Iron concentrations remained at baseline levels in both the fungal biomass and the agar regardless of lepidolite exposure. This is consistent with the crystal chemistry of lepidolite, in which Li⁺ substitutes for Al³⁺ within the octahedral sheet of the T-O-T mica structure, coordinated by bridging oxygens and fluorine, with negligible structural Fe (27, 28). This distinguishes lepidolite bioweathering from the bioleaching of many REE and critical mineral ores—such as lateritic deposits, Ce-bearing Fe-oxyhydroxides, or REE-adsorbed clays—where reductive dissolution of Fe(III) is a prerequisite for element liberation (2). The absence of Fe mobilization suggests that *Fusarium* sp. DS 682 accesses Li through direct proton- or ligand-promoted dissolution of the aluminosilicate framework rather than through Fe redox cycling, a mechanism consistent with the elevated organic acid production detected in our metabolomic analyses.

### Organic Acid Overflow as the Primary Mechanism of Lepidolite Dissolution

GC-MS metabolite profiling revealed that *lepidolite-grown* cultures secreted elevated concentrations of TCA cycle intermediates, particularly succinate, malate, and citrate, relative to uninoculated controls. Citric acid is a well-established biogenic lixiviant for aluminosilicate minerals due to its capacity to protonate mineral surfaces and to form stable complexes with Al³⁺ and other cations, and succinate and malate can also contribute to local pH depression and metal complexation (6, 7, 19).

Among the organic acids detected, oxalic acid was the only major species selectively elevated under lithium exposure relative to both uninoculated controls and sugar agar cultures. Oxalate is one of the most potent biogenic mineral-dissolving agents, capable of both proton-promoted dissolution of aluminosilicate frameworks and ligand-promoted dissolution through complexation of Al³⁺ and other structural cations (6, 18). Its selective upregulation on lepidolite—in contrast to citrate and succinate, which were elevated under both culture conditions—suggests that oxalate production represents a more specific metabolic response to lithium stress or mineral contact. Oxalic acid is synthesized via oxidative cleavage of oxaloacetate or through the glyoxylate cycle (6, 37), and its accumulation is consistent with the partially constrained TCA cycle inferred from the broader metabolite profiles.

Conversely, D-gluconic acid—a glucose oxidation product and itself a mineral-chelating organic acid—was markedly depleted in lepidolite-grown fungi compared to all other conditions. Gluconate is typically produced by extracellular glucose oxidase activity and can contribute to mineral dissolution through both acidification and Al/Fe complexation (1). Its depletion under lithium stress indicates that glucose carbon is redirected away from extracellular oxidation and toward intracellular pathways—specifically the INO1-mediated inositol rescues response, the pentose phosphate pathway, and the TCA cycle —that generate the myo-inositol, organic acids, polyols, and NADPH required for stress defense. This reallocation may represent a metabolic triage in which the fungus prioritizes intracellular survival over extracellular glucose oxidation.

The pattern of metabolite accumulation suggests that organic acids arise from a partially constrained TCA cycle operating under high flux. Under lithium stress, central carbon metabolism remains active—presumably to meet elevated ATP demands for ion pumping and cellular maintenance—but appears to encounter bottlenecks, possibly at the level of the respiratory chain or specific dehydrogenases perturbed by Li⁺. This results in incomplete oxidation and extracellular accumulation of TCA intermediates. Similar “overflow” secretion of organic acids has been documented in *Aspergillus niger* under nutrient limitation (6, 45); here, a comparable metabolic phenotype appears to be triggered by ionic rather than carbon or nitrogen stress.

Importantly, our data do not indicate that organic acid production is a targeted strategy evolved specifically to dissolve lepidolite. Instead, mineral dissolution emerges as a collateral consequence of the metabolic adjustments that allow the fungus to survive in a lithium-rich environment. This distinction has practical implications: it suggests that conditions promoting moderate, sub-lethal stress—rather than maximal growth—may yield the highest rates of organic acid secretion and, consequently, mineral dissolution.

### Metabolic Reprogramming in presence of Li: Growth Mode Versus Stress Mode

*Fusarium* sp. DS 682 reconfigures its metabolic program substantially in response to lepidolite. On sugar agar, the fungus operated in a growth-optimized mode characterized by vigorous glycolytic and TCA flux, high ergosterol, elevated unsaturated fatty acids supporting membrane fluidity, and active branched-chain amino acid (BCAA) catabolism. On lepidolite-amended medium, this program was reorganized toward stress defense: ergosterol and membrane fatty acids were suppressed, indicating constrained proliferation, while BCAA flux shifted from catabolism to biosynthesis (selective elevation of 2,3-dihydroxyisovalerate), suggesting prioritization of de novo amino acid synthesis to replace lithium-damaged enzymes. Concurrently, the glutamate pool expanded by more than an order of magnitude and L-serine – a precursor for cysteine via the transsulfuration pathway-was selectively elevated, consistent with increased demand for the rate-limiting substrate of glutathione biosynthesis. The dichotomy represents a classic growth-defense tradeoff: under lithium stress, the fungus sacrifices proliferative capacity to maintain ionic homeostasis. For bioleaching, this implies that while biomass-normalized acid production may be higher under stress, total leaching capacity could be constrained by reduced biomass, so optimizing ore concentration, carbon availability, and salinity will be critical.

### Compatible Solute Diversification and PPP-Linked Redox Defense

The accumulation of compatible solutes in lithium-stressed cultures is consistent with activation of ionic and osmotic stress responses, such as the high-osmolarity glycerol (HOG) pathway (46). Glycerol and trehalose were elevated under both conditions but on lepidolite-amended medium, the osmolyte pool expanded to include substantial amounts of erythritol and D-ribitol—polyols derived from pentose phosphate pathway (PPP) intermediates. This diversification suggests that ionic stress from Li⁺ presents a qualitatively different challenge than purely osmotic stress, requiring Non-reducing polyols that stabilize proteins and membranes without engaging in metal-catalyzed Maillard-type reactions (47). Because osmolyte production diverts glucose-derived carbon and, via glycerol formation, regenerates NAD⁺, it simultaneously supports osmotic balance and redox cofactor recycling. As this response competes with acid overflow for carbon, managing osmotic load offers a lever to redirect flux toward dissolution.

### PPP Upregulation Links Redox Defense, Osmolyte Production, and Mineral Chelation

The elevated D-ribitol and erythritol, together with lower pyruvate despite active TCA metabolism, indicate substantial PPP engagement under lithium stress.

The oxidative PPP generates NADPH required to maintain the glutathione redox cycle. Glutathione reductase consumes NADPH to regenerate reduced glutathione (GSH) from its oxidized form (GSSG), and the elevated L-pyroglutamic acid together with the expanded glutamate pool in lepidolite cultures indicate that this cycle operates at high capacity under lithium stress. In this sense, the PPP serves as a metabolic bridge between carbon metabolism and oxidative stress defense—an especially important connection under ionic stress, where both energetic and redox demands are simultaneously elevated. Because the PPP sustains both antioxidant defense and chelating polyols, conditions that support it may be key to maintaining long-term bioleaching performance.

### Oxidative Stress and Glutathione Cycling Under Lithium Exposure

Elevated L-pyroglutamic acid and 2-hydroxyglutaric acid in lithium-stressed cultures provide direct evidence of oxidative and redox imbalance. L-Pyroglutamic acid is the cyclized product of glutamate and accumulates when the γ-glutamyl cycle—which includes glutathione synthesis, utilization, and recycling—is operating at high flux (38). Its elevation, together with the greater than 30-fold expansion of the glutamate pool relative to uninoculated lepidolite controls, suggests active synthesis and turnover of glutathione to neutralize ROS generated under lithium stress. 2-Hydroxyglutaric acid, a recognized marker of mitochondrial dysfunction formed via reduction of α-ketoglutarate under high NADH/NAD⁺ ratios, co-occurred with expanded glutamate and pyroglutamate pools on lepidolite signaling strong demand on mitochondrial and glutathione-based defenses(40).

This oxidative stress response may have indirect implications for bioleaching. ROS can damage cellular membranes and proteins, potentially leading to cell lysis and release of intracellular metabolites, including organic acids, into the medium. Moreover, alternative pathways such as the GABA shunt, which converts glutamate to succinate via γ-aminobutyrate and succinate semialdehyde, may be activated under these conditions and contribute additional organic acids (e.g., 4-hydroxybutyrate) that further influence the mineral weathering environment. This links redox tolerance to leaching capacity: the longer the fungus sustains glutathione defense, the longer it stays active, making stress tolerance a useful strain-selection criterion.

### A Self-Reinforcing Cycle Links Stress Response to Mineral Dissolution

Synthesizing the ICP-MS, ToF-SIMS, morphological, and metabolite data, we propose a feedback model in which lithium bioleaching by *Fusarium* sp. DS 682 is coupled to the fungal stress response (Fig. 3). In the initial phase, fungal hyphae colonize the lepidolite surface, and constitutive organic acid secretion begins to dissolve the mineral at the hypha–mineral interface. As Li⁺ is liberated from the aluminosilicate lattice, it enters the fungal cell through K⁺ and Na⁺ transport systems that do not effectively discriminate Li⁺ from physiological monovalent cations (48). Once inside the cell, Li⁺ is poorly effluxed compared with Na⁺ and K⁺, leading to progressive intracellular Li⁺ loading. Because Li⁺ does not functionally replace K⁺ in most enzymes and ribosomes, this accumulation perturbs ionic balance, osmotic pressure, and mitochondrial function—as directly evidenced by the disruption of inositol phosphate metabolism, the activation of glutathione cycling, and the accumulation of the mitochondrial stress marker 2-hydroxyglutarate.

The resulting stress activates multiple defense pathways—upregulated central carbon metabolism for ATP generation, increased PPP flux for NADPH production and polyol synthesis, glutathione cycling for ROS scavenging, and naphthoquinone pigment overproduction for extracellular redox defense—all of which generate organic acid and polyol “overflow” products. These metabolites acidify and chelate the mineral surface, thereby promoting further Li⁺ release and reinforcing the cycle. Simultaneously, lithium-induced disruption of PIP₂-dependent polarity signaling drives morphological reorganization into hyphal cords, which concentrate organic acid secretion at the mineral interface and further enhance localized dissolution (49).

Whether this feedback is ultimately self-sustaining or self-limiting likely depends on the balance between the rate of Li⁺ release and the fungus’s capacity to tolerate intracellular Li⁺ loading. The observation that *Fusarium* sp. DS 682 maintained active metabolism and produced substantial biomass over the six-week incubation period suggests that the organism reaches a stress-adapted steady state rather than experiencing catastrophic Li⁺ poisoning. This contrasts with organisms that lack robust cation homeostasis, for which progressive Li⁺ loading would be expected to become lethal and terminate the bioleaching process. Analogous positive feedback loops have been described in bacterial bioleaching systems, where acid production by *Acidithiobacillus* spp. accelerates sulfide mineral oxidation, releasing additional substrates that fuel further bacterial growth and acid generation (9, 50). The fungal system described here represents a conceptually parallel mechanism driven predominantly by organic rather than sulfuric acids, with the additional complexity that morphological reorganization and secondary metabolite production contribute to the dissolution process.

## MATERIALS and METHODS

### Fungal Cultivation and Growth Conditions

*Fusarium* sp. DS 682, previously isolated from the rhizosphere of *Bouteloua gracilis* at the Konza Prairie Biological Station (KPBS) in Kansas (33), was maintained on a solid medium containing 0.1 M NaCl, 0.4% (w/v) glucose, and 0.1% (w/v) yeast extract (Difco 212750), solidified with 1.5% (w/v) agar (51). For bioleaching experiments, lepidolite (Standard Reference Material [SRM #183], National Institute of Standards and Technology [NIST], Gaithersburg, MD, USA) was incorporated into the medium at a concentration of 10% (w/v) prior to autoclaving.

### Bioleaching Assay

*Inoculation and incubation.* Three hyphal plugs (1.5 mm diameter) were excised from the margin of an actively growing colony using a sterile cork borer and transferred onto the surface of a sterile Sterlitech polyester (PETE) membrane overlaid on solidified medium with or without 10% (w/v) lepidolite. Inoculated plates were incubated at 25°C for six weeks. Three biological replicates were prepared per treatment.

*Fungal biomass and intracellular lithium quantification.* Following incubation, fungal biomass from three replicate plates per treatment was harvested by removing the membrane and associated mycelium, which were then lyophilized to constant weight. Dried mycelium was subsequently ground using a sterile mortar and pestle, and the resulting powder was extracted in 1 mL of 1 M sodium acetate (pH 8.2) by shaking at 220 rpm for 1 h at room temperature (∼22°C). Extracts were centrifuged at 7,500 rpm 10 min, and supernatants were collected for lithium quantification by inductively coupled plasma mass spectrometry (ICP-MS).

*Surface chemical analysis.* From an additional set of three replicate plates per treatment, mycelium was carefully detached from the membrane surface and transferred onto conductive copper tape for time-of-flight secondary ion mass spectrometry (ToF-SIMS) analysis.

*Agar extraction.* After fungal removal, the residual agar from each plate was bisected. One half was extracted with 1 M sodium acetate (pH 8.2) and the other with 70% (v/v) acetonitrile. Extractions were performed by shaking at 220 rpm for 1 h at room temperature (∼22°C). Samples were then centrifuged at 7,500 rpm for 10 min. Supernatants were collected and stored at -20°C until analysis by ICP-MS and gas chromatography–mass spectrometry (GC-MS), respectively.

### Inductively Coupled Plasma Mass Spectrometry (ICP-MS)

The sodium acetate extract samples were analyzed for Lithium using an Agilent 8900 triple quadrupole ICP-MS (Agilent Technologies, Inc., Santa Clara, CA, USA). Prior to analysis, extracts were diluted 10-fold in 2% (v/v) nitric acid (trace-metal grade). Method blanks consisting of sterile sodium acetate extraction buffer processed with uninoculated PETE membranes were diluted and analyzed identically.

### Time-of-Flight Secondary Ion Mass Spectrometry (ToF-SIMS)

ToF-SIMS analysis was performed at the Environmental Molecular Sciences Laboratory (EMSL) at Pacific Northwest National Laboratory (Richland, WA, USA) using a TOF.SIMS 5 instrument (IONTOF GmbH, Münster, Germany). Data were acquired in both high-mass-resolution spectral mode and high-spatial-resolution imaging mode in positive ion polarity.

*High-mass-resolution spectral mode.* A 25 keV Bi⁺ primary ion beam was focused to an approximately 5 µm spot diameter with a beam current of 1.69 pA at a 10 kHz pulse repetition frequency. Spectra were collected over a 300 × 300 µm² field of view for 20 scans. Under these conditions, mass resolving power (*m*/Δ*m*) was typically in the range of 4,000–8,000.

*High-spatial-resolution imaging mode.* The Bi⁺ beam was focused to an approximately 400 nm spot diameter and rastered over a 100 × 100 µm² field of view at 256 × 256 pixels with a cycle time of 50 µs, yielding unit-mass resolution in the corresponding spectra.

For both acquisition modes, charge compensation was achieved using an electron flood gun (∼14.17 µA). Ion spectra were processed and extracted using SurfaceLab software (version 7.2; IONTOF GmbH).

### Untargeted metabolomics analysis by GC-MS

Extracted metabolites were chemically derivatized and analyzed as untargeted manner by gas chromatography-mass spectrometer (GC-MS) as reported previously (52). Briefly, extracted metabolites after derivatization were injected into GC-MS and their chromatographic separations and MS fragmentation patterns were matched against PNNL in-house databases originally derived from Agilent Fiehn Metabolomics database. In addition, other public and commercially available databases (NIST23, Wiley 11^th^ edition) were used to cross-check metabolite identification. Identification of metabolites and their peak area values were manually curated when needed.

## ACKNOWLEDGEMENTS

This research was supported by the U.S. Department of Energy (DOE), Office of Biological and Environmental Research (OBER), as part of BER’s Genomic Science Program (GSP), under project 82324 “Terraforming Soil EERC: Accelerating Soil-Based Carbon Drawdown through Advanced Genomics and Geochemistry.” A portion of this work was performed in the William R. Wiley Environmental Molecular Sciences Laboratory (EMSL), a national scientific user facility sponsored by OBER and located at PNNL. PNNL is a multi-program national laboratory operated by Battelle for the DOE under Contract DE-AC05-76RLO 1830.

## Reference

1. Gadd GM. 2007. Geomycology: biogeochemical transformations of rocks, minerals, metals and radionuclides by fungi, bioweathering and bioremediation. Mycological Research 111:3– 49.

2. Gadd GM. 2010. Metals, minerals and microbes: geomicrobiology and bioremediation. Microbiology 156:609–643.

3. Burford EP, Fomina M, Gadd GM. 2003. Fungal involvement in bioweathering and biotransformation of rocks and minerals. Mineralogical Magazine 67:1127–1155.

4. Uroz S, Calvaruso C, Turpault M-P, Frey-Klett P. 2009. Mineral weathering by bacteria: ecology, actors and mechanisms. Trends in Microbiology 17:378–387.

5. Aishvarya V, Barman S, Pradhan N, Ghosh MK. 2019. Selective enhancement of Mn bioleaching from ferromanganese ores in presence of electron shuttles using dissimilatory Mn reducing consortia. Hydrometallurgy 186:269–274.

6. Gadd GM. 1999. Fungal Production of Citric and Oxalic Acid: Importance in Metal Speciation, Physiology and Biogeochemical Processes, p. 47–92. In Advances in Microbial Physiology. Academic Press.

7. Rezza I, Salinas E, Elorza M, Sanz de Tosetti M, Donati E. 2001. Mechanisms involved in bioleaching of an aluminosilicate by heterotrophic microorganisms. Process Biochemistry 36:495–500.

8. Ahmed E, Holmström SJM. Siderophores in environmental research:roles and applications.

9. Song X, Yang A, Hu X, Niu A, Cao Y, Zhang Q. 2023. Exploring the role of extracellular polymeric substances in the antimony leaching of tailings by Acidithiobacillus ferrooxidans. Environ Sci Pollut Res 30:17695–17708.

10. Ye M, Liang J, Liao X, Li L, Feng X, Qian W, Zhou S, Sun S. 2021. Bioleaching for detoxification of waste flotation tailings: Relationship between EPS substances and bioleaching behavior. Journal of Environmental Management 279:111795.

11. Drever JI, Stillings LL. 1997. The role of organic acids in mineral weathering. Colloids and Surfaces A: Physicochemical and Engineering Aspects 120:167–181.

12. Kraemer SM, Crowley DE, Kretzschmar R. 2006. Geochemical Aspects of Phytosiderophore-Promoted Iron Acquisition by Plants, p. 1–46. In Advances in Agronomy. Academic Press.

13. Saha R, Saha N, Donofrio RS, Bestervelt LL. 2013. Microbial siderophores: a mini review. Journal of Basic Microbiology 53:303–317.

14. Amiri F, Yaghmaei S, Mousavi SM, Sheibani S. 2011. Recovery of metals from spent refinery hydrocracking catalyst using adapted Aspergillus niger. Hydrometallurgy 109:65–71.

15. Horeh NB, Mousavi SM, Shojaosadati SA. 2016. Bioleaching of valuable metals from spent lithium-ion mobile phone batteries using Aspergillus niger. Journal of Power Sources 320:257–266.

16. Moazzam P, Boroumand Y, Rabiei P, Baghbaderani SS, Mokarian P, Mohagheghian F, Mohammed LJ, Razmjou A. 2021. Lithium bioleaching: An emerging approach for the recovery of Li from spent lithium ion batteries. Chemosphere 277:130196.

17. Sedlakova-Kadukova J, Marcincakova R, Luptakova A, Vojtko M, Fujda M, Pristas P. 2020. Comparison of three different bioleaching systems for Li recovery from lepidolite. Sci Rep 10:14594.

18. Fomina M, Hillier S, Charnock J, Melville K, Alexander I, Gadd G. 2005. Role of Oxalic Acid Overexcretion in Transformations of Toxic Metal Minerals by Beauveria caledonica. Applied and environmental microbiology 71:371–81.

19. Wei Z, Kierans M, Gadd GM. 2012. A Model Sheet Mineral System to Study Fungal Bioweathering of Mica. Geomicrobiology Journal 29:323–331.

20. Yongdong W, Wang J, Dexin, Ding, Li G, Sun J, hu N, Li F, Ma J, Zhang H, Ding Y, Dai Z. 2023. Hyphae and organic acids of Aspergillus Niger promote uranium recovery by destroying the ore surface and increasing the porosity and permeability of ores. Nuclear Engineering and Technology 10.1016/j.net.2023.12.049.

21. Dusengemungu L, Kasali G, Gwanama C, Mubemba B. 2021. Overview of fungal bioleaching of metals. Environmental Advances 5:100083.

22. Dong Y, Zan J, Lin H. 2023. Bioleaching of heavy metals from metal tailings utilizing bacteria and fungi: Mechanisms, strengthen measures, and development prospect. Journal of Environmental Management 344:118511.

23. Owusu-Fordjour EY, Yang X. 2023. Bioleaching of rare earth elements challenges and opportunities: A critical review. Journal of Environmental Chemical Engineering 11:110413.

24. Guo H, Kuang G, Wan H, Yang Y, Yu H, Wang H. 2019. Enhanced acid treatment to extract lithium from lepidolite with a fluorine-based chemical method. Hydrometallurgy 183:9–19.

25. Duan H, Zhao X, Xu C, Zhang D, Gu W, Wang R, Lu X. 2025. The synergistic interaction of fungi with different structural characteristics improves the leaching of lithium from lepidolite. Biochemical Engineering Journal 213:109564.

26. Tian-ming G, Na F, Wu C, Tao D, MNR Key Laboratory of Metallogeny and Mineral Assessment, Institute of Mineral Resources Chinese Academy of Geological Sciences, Beijing 100037, China, Research Center for Strategy of Global Mineral Resources, Chinese Geological Survey, Beijing 100037, China, China Huanqiu Contracting & Engineering Corp., LTD Beijing Branch, Beijing 100012, China, SDU Life Cycle Engineering, Department of Green Technology, University of Southern Denmark, Odense 5230, Denmark. 2023. Lithium extraction from hard rock lithium ores (spodumene, lepidolite, zinnwaldite, petalite): Technology, resources, environment and cost. China Geology 6:137–153.

27. Foster MD. 1960. Interpretation of the composition of trioctahedral micas. 354-B. Professional Paper 10.3133/pp354B.

28. Deer WA, Howie RA, Zussman J. 2013. An Introduction to the Rock-Forming Minerals. Mineralogical Society of Great Britain and Ireland. https://pubs.geoscienceworld.org/minersoc/books/book/952/An-Introduction-to-the-Rock-Forming-Minerals. Retrieved 23 April 2026.

29. Hallcher LM, Sherman WR. 1980. The effects of lithium ion and other agents on the activity of myo-inositol-1-phosphatase from bovine brain. Journal of Biological Chemistry 255:10896–10901.

30. Jesch SA, Zhao X, Wells MT, Henry SA. 2005. Genome-wide analysis reveals inositol, not choline, as the major effector of Ino2p-Ino4p and unfolded protein response target gene expression in yeast. J Biol Chem 280:9106–9118.

31. Shirra MK, Patton-Vogt J, Ulrich A, Liuta-Tehlivets O, Kohlwein SD, Henry SA, Arndt KM. 2001. Inhibition of Acetyl Coenzyme A Carboxylase Activity Restores Expression of the INO1 Gene in a snf1 Mutant Strain of Saccharomyces cerevisiae. Mol Cell Biol 21:5710–5722.

32. Vaden DL, Ding D, Peterson B, Greenberg ML. 2001. Lithium and valproate decrease inositol mass and increase expression of the yeast INO1 and INO2 genes for inositol biosynthesis. J Biol Chem 276:15466–15471.

33. Bhattacharjee A, Anderson LN, Alfaro T, Porras-Alfaro A, Jumpponen A, Hofmockel KS, Jansson JK, Anderton CR, Nelson WC. 2021. Draft Genome Sequence of Fusarium sp. Strain DS 682, a Novel Fungal Isolate from the Grass Rhizosphere. Microbiol Resour Announc 10:e00884–20.

34. Boddy L. 1999. Saprotrophic cord-forming fungi: meeting the challenge of heterogeneous environments. Mycologia 91:13–32.

35. Lindahl BD, Olsson S. 2004. Fungal translocation – creating and responding to environmental heterogeneity. Mycologist 18:79–88.

36. Cairney JWG. 1992. Translocation of solutes in ectomycorrhizal and saprotrophic rhizomorphs. Mycological Research 96:135–141.

37. Munir E, Yoon JJ, Tokimatsu T, Hattori T, Shimada M. 2001. A physiological role for oxalic acid biosynthesis in the wood-rotting basidiomycete Fomitopsis palustris. Proceedings of the National Academy of Sciences 98:11126–11130.

38. Pócsi I, Prade R, Penninckx M. 2004. Glutathione, Altruistic Metabolite in Fungi. Advances in microbial physiology 49:1–76.

39. Thomas D, Surdin-Kerjan Y. 1997. Metabolism of sulfur amino acids in Saccharomyces cerevisiae. Microbiol Mol Biol Rev 61:503–532.

40. Intlekofer AM, Dematteo RG, Venneti S, Finley LWS, Lu C, Judkins AR, Rustenburg AS, Grinaway PB, Chodera JD, Cross JR, Thompson CB. 2015. Hypoxia Induces Production of L-2-Hydroxyglutarate. Cell Metab 22:304–311.

41. Guillén DR, Gimenes LJ, Romano Espinosa DC, Soares Tenório JA, Baltazar M dos PG. 2026. Fungal biosorption in lithium-ion battery recycling: A review of critical metal recovery. Journal of Environmental Management 401:128892.

42. Studt L, Wiemann P, Kleigrewe K, Humpf HU, Tudzynski B. 2012. Biosynthesis of fusarubins accounts for pigmentation of fusarium Fujikuroi perithecia. Applied and Environmental Microbiology 78:4468–4480.

43. Frandsen RJN, Nielsen NJ, Maolanon N, Sørensen JC, Olsson S, Nielsen J, Giese H. 2006. The biosynthetic pathway for aurofusarin in Fusarium graminearum reveals a close link between the naphthoquinones and naphthopyrones. Molecular Microbiology 61:1069–1080.

44. Malz S, Grell MN, Thrane C, Maier FJ, Rosager P, Felk A, Albertsen KS, Salomon S, Bohn L, Schäfer W, Giese H. 2005. Identification of a gene cluster responsible for the biosynthesis of aurofusarin in the *Fusarium graminearum* species complex. Fungal Genetics and Biology 42:420–433.

45. Papagianni M. 2007. Advances in citric acid fermentation by *Aspergillus niger*: Biochemical aspects, membrane transport and modeling. Biotechnology Advances 25:244–263.

46. Hohmann S. 2002. Osmotic Stress Signaling and Osmoadaptation in Yeasts. Microbiology and Molecular Biology Reviews 66:300–372.

47. Yancey PH. 2005. Organic osmolytes as compatible, metabolic and counteracting cytoprotectants in high osmolarity and other stresses. J Exp Biol 208:2819–2830.

48. Rodrýguez-Navarro A. 2000. Potassium transport in fungi and plants. Biochimica et Biophysica Acta (BBA) - Reviews on Biomembranes 1469:1–30.

49. Schultzhaus ZS, Shaw BD. 2015. Endocytosis and exocytosis in hyphal growth. Fungal Biology Reviews 29:43–53.

50. Zhang S, Yang J, Dong B, Yang J, Pan H, Wang W, Yan L, Gu J-D. 2022. An Fe(II)-oxidizing consortium from Wudalianchi volcano spring in Northeast China for bioleaching of Cu and Ni from printed circuit boards (PCBs) with the dominance of Acidithiobacillus spp. International Biodeterioration & Biodegradation 167:105355.

51. Song W, Ogawa N, Oguchi CT, Hatta T, Matsukura Y. 2007. Effect of *Bacillus subtilis* on granite weathering: A laboratory experiment. CATENA 70:275–281.

52. King WL, Yates CF, Cao L, O’Rourke-Ibach S, Fleishman SM, Richards SC, Centinari M, Hafner BD, Goebel M, Bauerle T, Kim Y-M, Nicora CD, Anderton CR, Eissenstat DM, Bell TH. 2023. Functionally discrete fine roots differ in microbial assembly, microbial functional potential, and produced metabolites. Plant, Cell & Environment 46:3919–3932.

